## Supplemental information for "Temperature sensitivity of bat antibodies links metabolic state with antigen-recognition diversity"

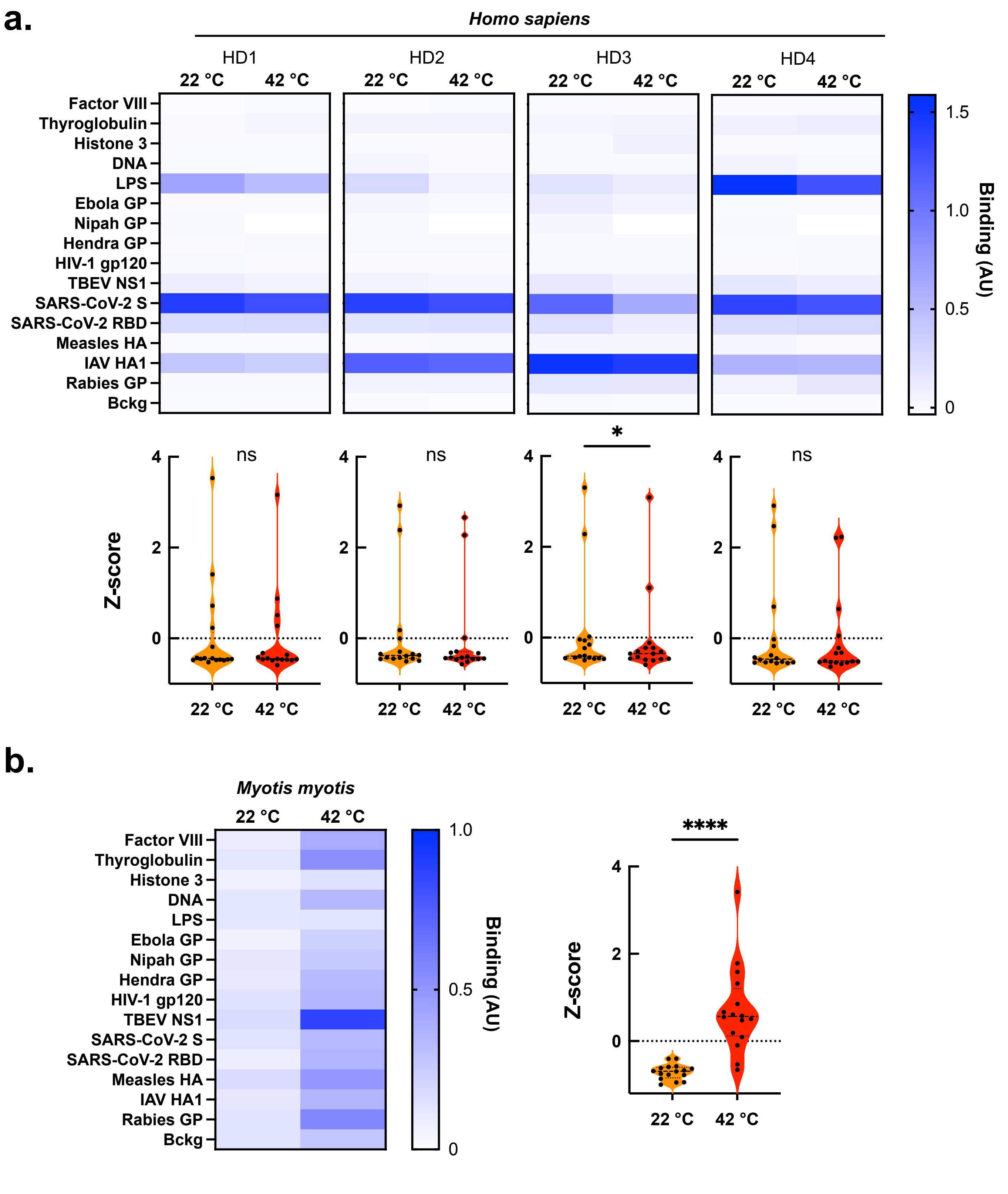


**Suppl. Figure 1.** Temperature differentially modulates the antigen binding properties of bat and human IgG. **a.** Heat maps depicting the binding intensity of IgG purified from sera of 4 healthy humans to a panel of antigens. **b**. Heat map depicting the reactivity of IgG purified from *M. myotis* pooled sera (n=5). The reactivity of bat and human IgG was evaluated at 22 and 42 °C by ELISA. The binding of IgG to the blocking agent (Bckg) is also shown. The binding intensity against each target represent average optical density (n=2) after subtraction of the background binding (measured after incubation of the detection reagents with the respective antigens in absence of antibodies). The violin plots corresponding to each heat map show the z-scores values of antigen binding. The z-score for each target antigen is presented as an individual circle. *P* values were determined using Wilcoxon matched-pairs signed rank test, ****$p < 0.0001$, n.s., not significant.

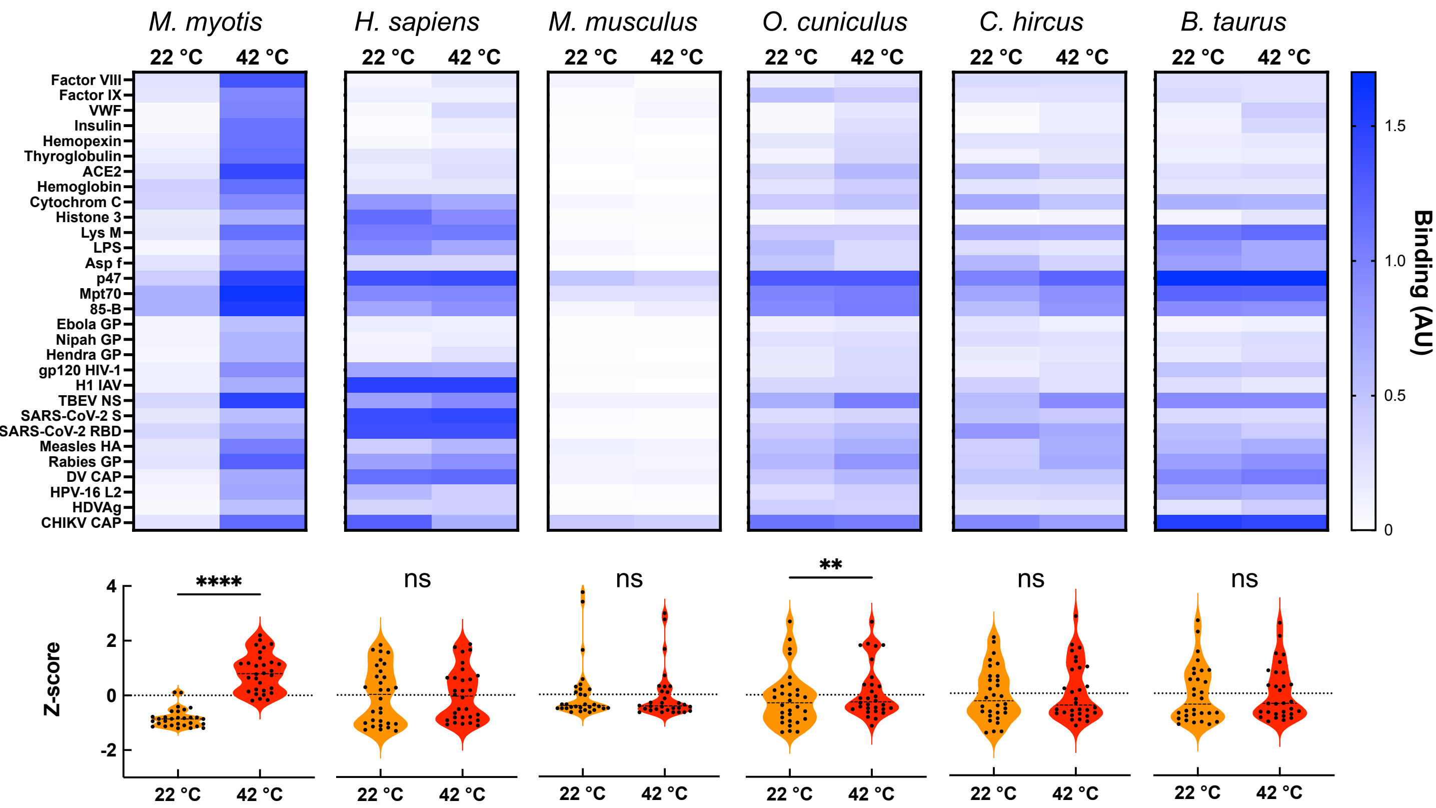


**Suppl. Figure 2.** Temperature has a differential effect on the antigen-binding activity of IgG from different mammal species. *Upper panels* Heat maps depicting the binding intensity of IgG purified from sera pools of bat (*Myotis myotis,* pool n=10 individuals), human (*Homo sapiens,* pool n=10 individuals), mouse (*Mus musculus*), rabbit (*Oryctolagus cuniculus*), goat (*Capra hircus*), and cattle (*Bos taurus*) to a panel of distinct antigens. The binding purified IgG was evaluated at 22 and 42 °C by ELISA. The binding intensity against each target represent average optical density (n=2) after subtraction of the background binding (measured after incubation of the detection reagents, protein G-biotin and streptaviding-HRP with the respective antigens in absence of IgG). *Lower panels* Violin plots corresponding to each heat map show the z-scores values of antigen binding. The z-score for each target antigen is presented as an individual circle. *P* values were determined using Wilcoxon matched-pairs signed rank test, ****$p < 0.0001$, **$p < 0.02$, n.s., not significant.

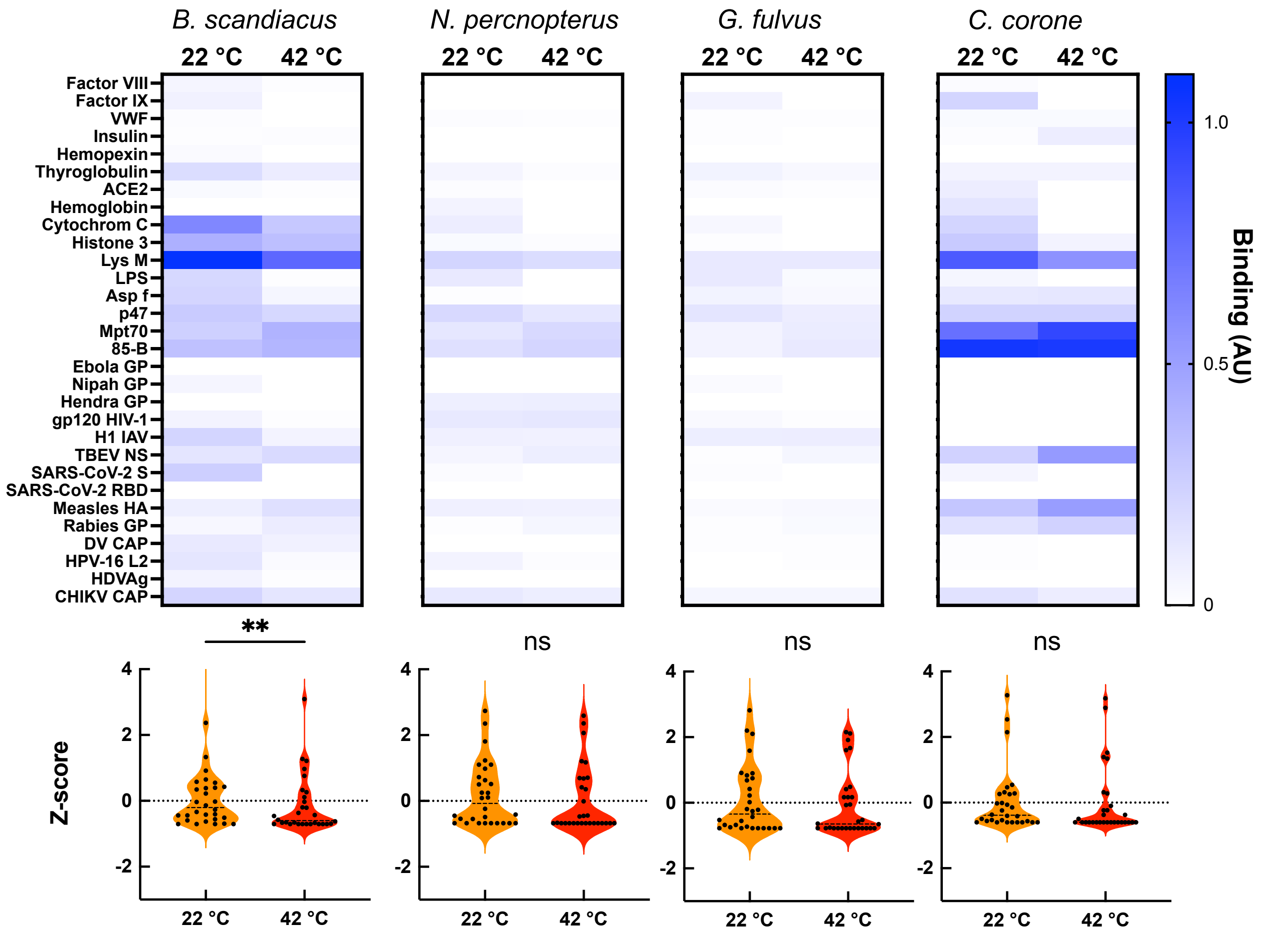


**Suppl. Figure 3.** Effect of temperature on the antigen binding properties of bird IgY antibodies. Heat maps depict the binding intensity of IgY in pooled sera of different bird species (*Bubo scandiacus*, n=2; *Neophron percnopterus,* n=2; *Gyps fulvus,* n=2, and Corvus corone, n=4) to a panel of antigens. The reactivity of bird IgY was evaluated at 22 and 42 °C by ELISA. The binding intensity, presented in arbitrary units, was obtained after subtraction of the reactivity of sera to the control surface (without coated antigen). In cases when the substation resulted in negative values the value of 0 was given. The violin plots corresponding to each heat map show the z-scores values of antigen binding by IgY. The z-score for each target antigen is presented as an individual circle. *P* values were determined using Wilcoxon matched-pairs signed rank test, $^{**}p < 0.01$, n.s., not significant.

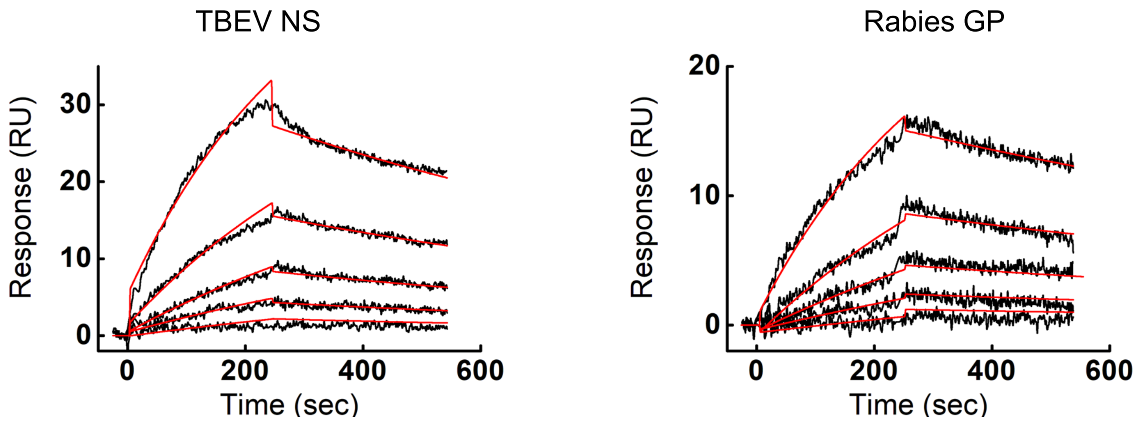


**Suppl. Figure 4.** Real-time interaction profiles showing the binding of IgG from *M. myotis* at 40 °C to immobilized on sensor chip Rabies virus glycoprotein and TBEV NS1 antigen. The antibodies were injected at serial dilution in concentration range 670 – 41.875 nM. The apparent binding affinities were evaluated by global Langmuir kinetics analyses model. The experimental data are shown by black lines. The kinetic fit is depicted by red lines.

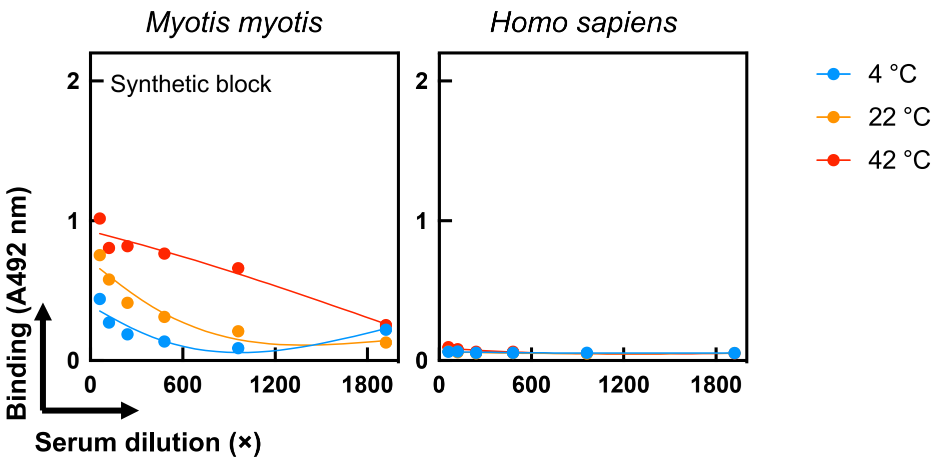


**Suppl. Figure 5.** ELISA assay for assessing the effect of temperature on binding to the blocked surface by bat and human IgG in whole sera. Pooled bat and human sera were serially diluted from 60 × to 1920 × and incubated with surface blocked by Synthetic Block (Thermo Fisher Scientific) at 4, 22 and 42 °C. Each data point represents average IgG binding intensity ±SD from n=3 technical replicates.

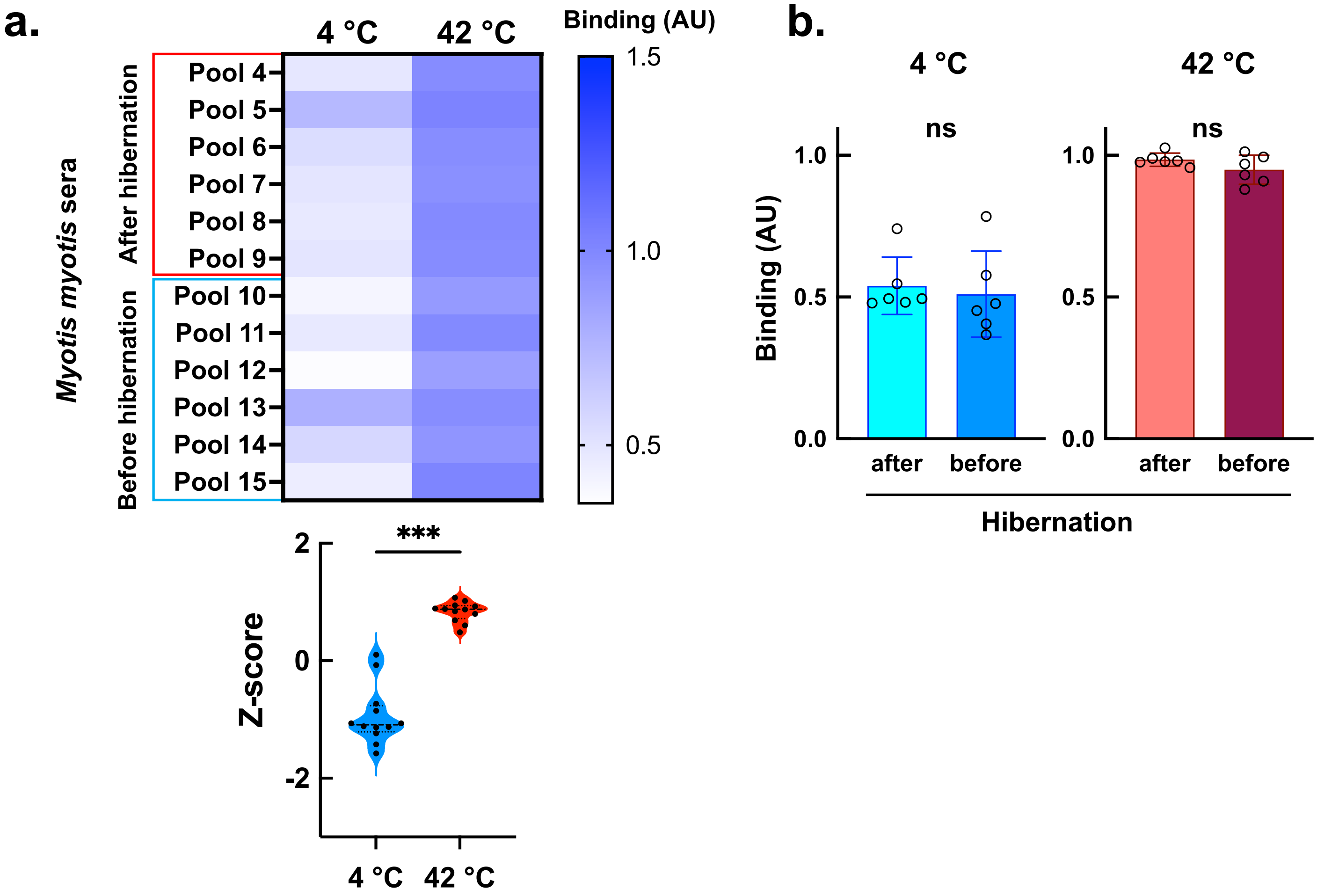


**Suppl. Figure 6.** Temperature uncovers antigen-binding specificity in different sera pools from *M. myotis*. **a.** Heat map depicting the reactivity of IgG to NS1 from TBEV in whole sera pools from *M. myotis* (n=3-8 individuals each). The sera were collected in different seasons – in spring (after hibernation) and in autumn (before hibernation) The reactivity of bat IgG in whole sera was evaluated at 4 and 42 °C by ELISA.. The binding intensity against NS1 of TBEV represent average optical density (n=2) after subtraction of the background binding (measured after incubation of the detection reagents with the respective antigens in absence of antibodies). The violin plots corresponding to each heat map show the z-scores values of antigen binding. The z-score for each target antigen is presented as an individual circle. *P* values were determined using Wilcoxon matched-pairs signed rank test, $^{***}p = 0.0005$. **b**. Bar graphs analyze the effect of time of collection of samples on the temperature sensitivity of binding of bat IgG in whole sera to NS1 of TBEV. Each circle represents individual serum pool. The statistical significance was analyzed by using nonparametric t-test, Mann-Whitney test, n.s. not significant.

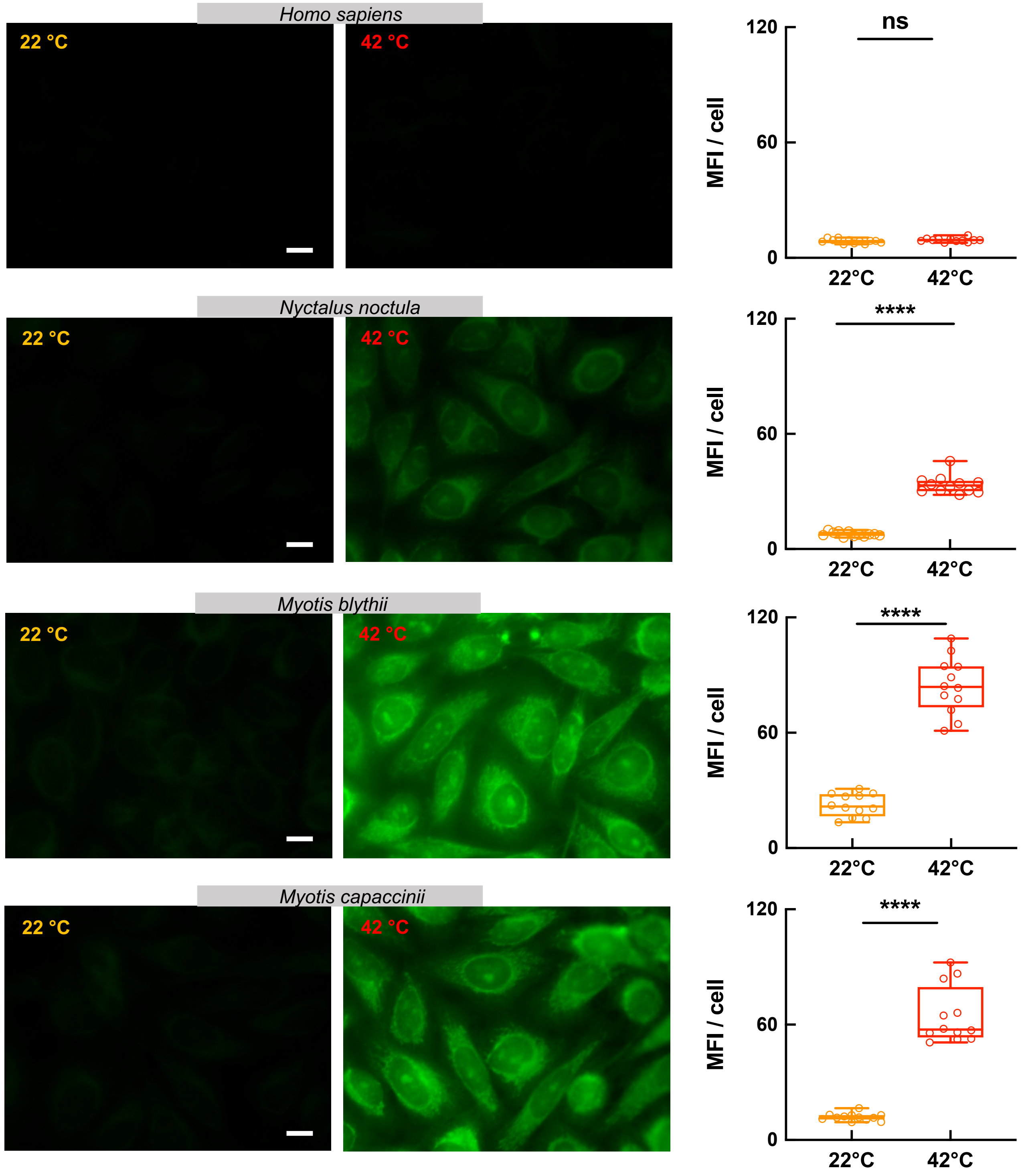


**Suppl. Figure 7.** Immunofluorescence analyses of binding of bat (*Nyctalus noctula*, *Myotis blythii*, and *Myotis capaccinii*) and human IgG to HEp-2 cells at 22 and 42 °C. The images are representative examples from n=12 fluorescence acquisitions from two independent experiments (magnification × 63). Right panels shows quantification of fluorescence intensity. The graphs show the mean fluorescence intensity of n=12 images, for each condition, acquired in two independent experiments. Statistical analyses were performed by using nonparametric t-test, Mann-Whitney test, $****p < 0.0001$. The white bar corresponds to 10 $\mu$m.

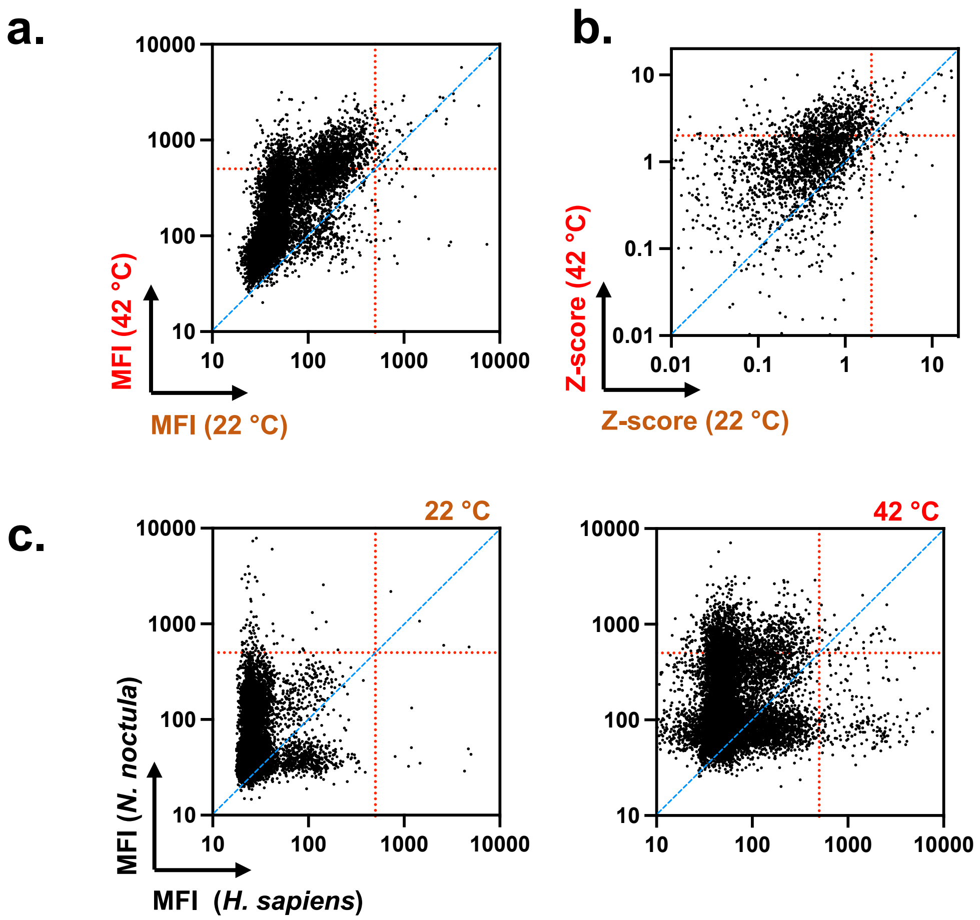


**Suppl. Figure 8.** IgG antibodies from *Nyctalus noctula* broaden their reactivity towards distinct proteins at elevated temperature. The presented plots summarize results from protein microarray analyses of binding of purified IgG antibodies from *N. noctula* to human proteins. In a. the mean fluorescence intensity (MFI) of pooled IgG reactivity at 22 °C was plotted versus the reactivity at 42 °C; in b. the Z-scores of antibody binding reactivity of *N. noctula* at 22 °C was plotted versus the reactivity at 42 °C. In c. the antibody reactivity (MFI) of human pooled IgG versus pooled IgG from *N. noctula* was plotted at 22 and 42 °C. Each dot represent MFI or Z-score signifying the binding of IgG to a single protein in the array, after substruction of the background reactivity. Red dashed lines represent an arbitrary threshold of 500 MFI. The threshold indicated with red dotted lines in b corresponds to Z=2 and $p < 0.05$. The data for human IgG are used for reference and are identical as those displayed on main Fig. 2.

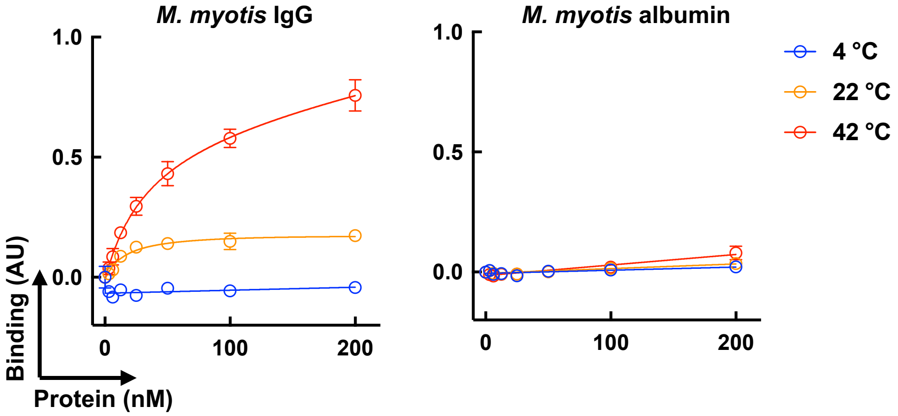


**Suppl. Figure 9.** ELISA assay for comparing the effect of temperature on binding of bat IgG and bat albumin to crude lysate of bat cells. Pooled bat IgG (*M. myotis*) and biotinylated albumin purified from pooled sera (n=10) of *M. myotis* were serially diluted from 200 to 3.125 nM and incubated with surfaces immobilized crude extract of epithelial cells (MmNep) from *M. myotis*. The proteins were incubated at 4, 22 and 42 °C. Each data point represents average IgG binding intensity ±SD (n=3 IgG and n=6 albumin, values combined from two independent experiments). The values were obtained after subtraction of background binding measured after incubation of the detection reagents with the coated wells in absence of antibodies or albumin.

**a.**

```
hIgG1                    ASTKGPSVFPLAPSSKSTSGGTAALGCLVKDYFPEPVTVSWNSGALTSGVHTFPAVLQSS   60
M.myo_NW_023416391.1     ASTTAPSVFPLSPKCGTTSSSTVSLGCLVSGYFPEPVTVTWNSGSLTSGVHTFPSVLR-S   59
M.myo_NW_023416348.1     ASTTAPSVFPLSPKCGTTSGSTVSLGCLVSGYFPEPVTVTWNSGSLTSGVHTFPSVLR-S   59
                         ***..******:*.. :**..*.:*****..********:****:*********:**: *

hIgG1                    GLYSLSSVVTVPSSSLGTQTYICNVNHKPSNTKVDKKVEPKSC-----DKTHTCPPCPAP   115
M.myo_NW_023416391.1     GLYSMSSMVTVPASS-STQTFICNVAHPASSTKVDKIVPLTVGPPPTSCPPHTCPPCE--   116
M.myo_NW_023416348.1     GLYSMSSMVTVPASS-STQTFICNVAHPASSTKVDKTVPLTVGPPPTSCPPHTCPPCE--   116
                         ****:**:****:** .***:**** *  *.***** *   .         ******

hIgG1                    ELLGGPSVFLFPPKPKDTLMISRTPEVTCVVVDVSHEDPEVKFNWYVDGVEVHNAKTKPR   175
M.myo_NW_023416391.1     -NPGGPSVFIFPPKPKDTLMISRTPEVTCMVVDVAPDDLDVEFTWYMDGNQMTKKTVKAE   175
M.myo_NW_023416348.1     -NPGGPSVFIFPPKPKDTLMISRTPEVTCMVVDVAPDDLDVEFTWYMDGNQMTKKTVKAE   175
                            ******:*******************:****: :* :*:*.**:** :: : ..* .

hIgG1                    EEQYNSTYRVVSVLTVLHQDWLNGKEYKCKVSNKALPAPIEKTISKAKGQPREPQVYTLP   235
M.myo_NW_023416391.1     QEQFNSTYRVVHSYAITHQDWLKGKKFKCKVNNKAIPSPIERTISKAIGQVQAPQVYVLG   235
M.myo_NW_023416348.1     QEQFNSTYRVVHSYAITHQDWLKGKKFKCKVNNKAIPSPIERTISKAIGQVQAPQVYVLG   235
                         :**:*******   :: *****:**::****.***:*:***:***** ** : ****.*

hIgG1                    PSRDELTKNQVSLTCLVKGFYPSDIAVEWESNGQPEN--NYKTTPPVLDSDGSFFLYSKL   293
M.myo_NW_023416391.1     PHSDELARDKVSVTCLVKDFFPPDISVEWQSNGQPESETKYSSTPPQKDQEGSFFLYSKL   295
M.myo_NW_023416348.1     PHSDELARDKVSVTCLVKDFFPPDISVEWQSNGQPESETKYSSTPPQKDQEGSFFLYSKL   295
                         *  ***::::**:*****.*:* **:***:******.  :*.:***  *.:*********

hIgG1                    TVDKSRWQQGNVFSCSVMHEALHNHYTQKSLSLSPGK        330
M.myo_NW_023416391.1     TVDKARWQRGAPFTCEVMHEGLHNHYAQKTVSWNPGK        332
M.myo_NW_023416348.1     TVDKARWQRGAPFTCEVMHEGLHNHYAQKTVSWNPGK        332
                         ****:***:*  *:*.****.*****:**::* .***
```

- CH1 Domain
- Hinge
- CH2 Domain
- CH3 Domain

**b.**

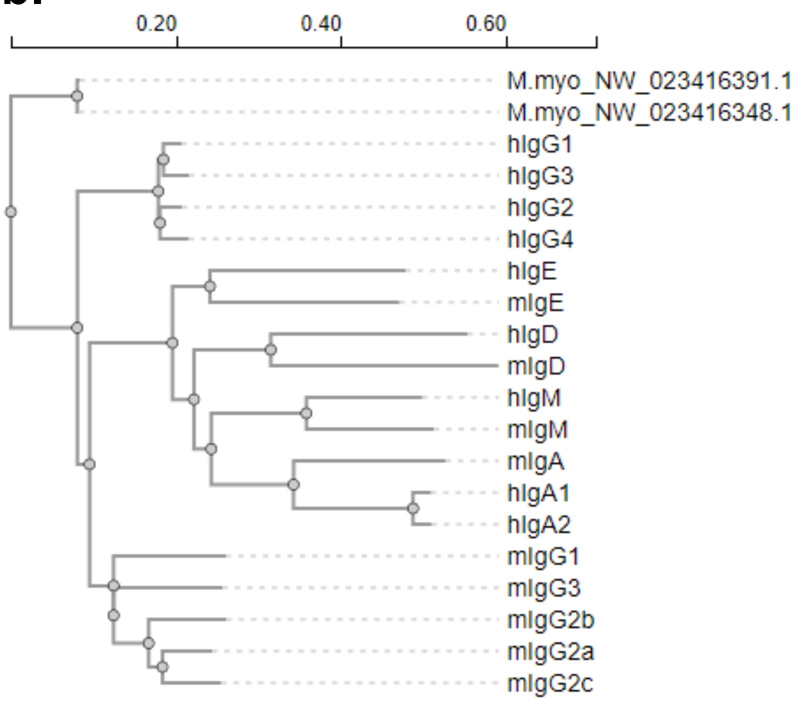


**c.**

| Sequence | Percent Identity |
|---|---|
| hIgG1 | 67,13 |
| hIgG2 | 64,535 |
| hIgG4 | 64,17 |
| hIgG3 | 62,88 |
| mIgG2b | 58,435 |
| mIgG3 | 58,415 |
| mIgG2a | 58,23 |
| mIgG2c | 58,13 |
| mIgG1 | 57,68 |
| hIgE | 36,09 |
| mIgE | 35,35 |
| hIgM | 31,03 |
| hIgA1 | 30,79 |
| hIgA2 | 30,46 |
| mIgM | 29,69 |
| mIgA | 28,15 |
| hIgD | 25,24 |
| mIgD | 20,27 |

**Suppl. Figure 10.** Sequence analyses of Fc-portion of bat IgG. **a.** Alignment of the sequences of human Fc-$\gamma$1 chain with two alotypes (NW_023416391.1 and NW_023416348.1) of Fc-$\gamma$1 chain of Myotis myotis. The different domains in the constant regions are indicated by colors. **b.** Phylogenetic tree depicting the homology of bat Fc-$\gamma$1 chains with constant chains of different classes and subclasses of heavy chains of human and mouse immunoglobulins. **c**. Percentage identity in the sequence of Fc-$\gamma$1 chain of *M. myotis* to different human and mouse immunoglobulin heavy chains.

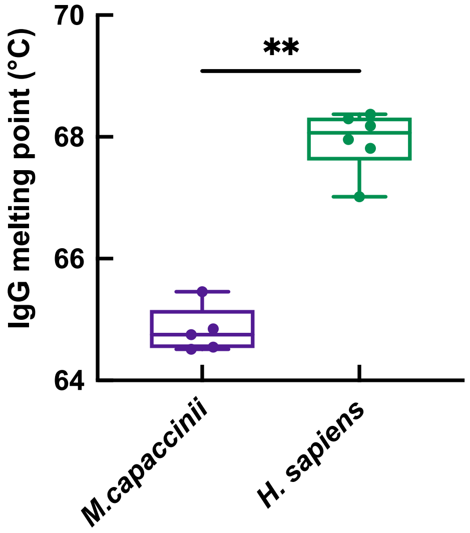


**Suppl. Figure 11.** Comparison of the thermodynamic stability of IgG from bat and human. Graph depicting box plot of melting temperatures of bat (*M. capaccinii*) and human IgG, obtained by thermal-shift assay. Each dot in the plots indicate individual estimation of the melting temperature. The results of two independent experiments are presented together. The statistical significance was assessed by applying nonparametric t-test analyses, Mann-Whitney test. **$p = 0.0043$.

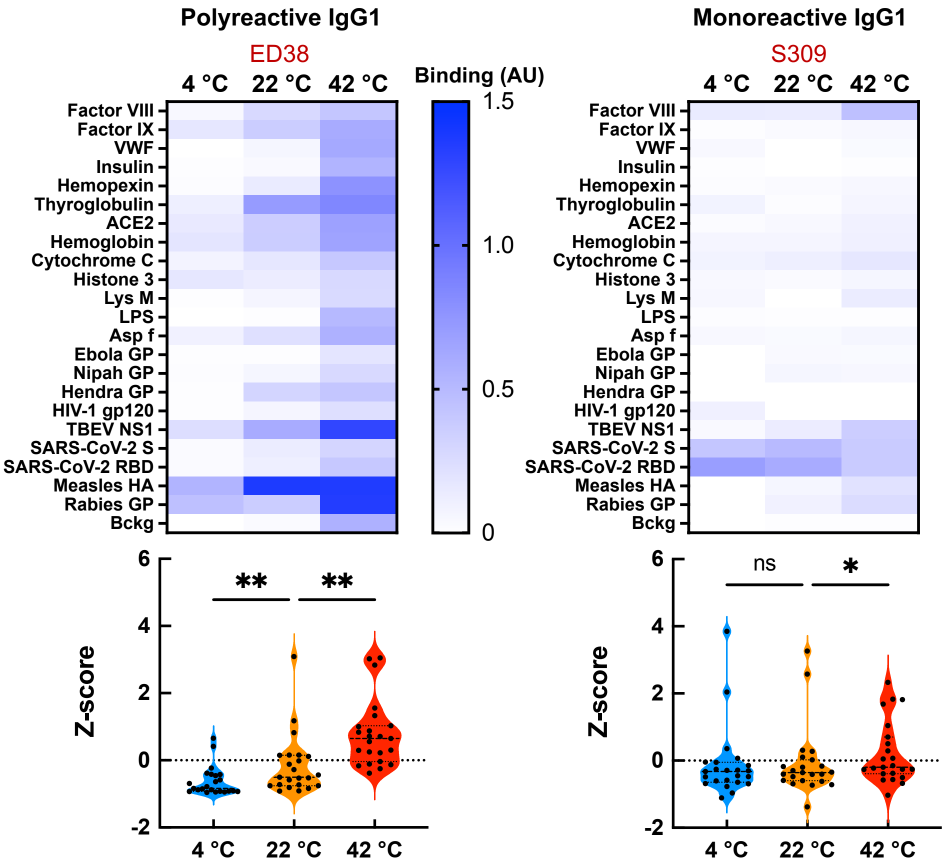


**Suppl. Figure 12.** Temperature-dependence of antigen binding by polyreactive and monoreactive human IgG antibodies. Heat map depicting the binding intensity of human polyreactive (ED38) and monoreactive (S309) IgG1 to a panel of distinct antigens. The reactivity of monoclonal antibodies was evaluated at 4, 22 and 42 °C by ELISA. The binding of IgG to the blocking agent (Bckg) is also shown. The binding intensity against each target represent average optical density (n=2) after subtraction of the background binding (measured after incubation of the detection reagents to the respective antigens in absence of IgG). For analyses both antibodies were diluted to 30 µg/ml. The violin plots corresponding to each heat map show the Z-scores values of antigen binding. The Z-score for each target antigen is presented as an individual circle. P values were determined using one-way analysis of variance (ANOVA) with Friedman multiple comparison test, *$p < 0.02$, **$p < 0.005$, n.s., not significant.

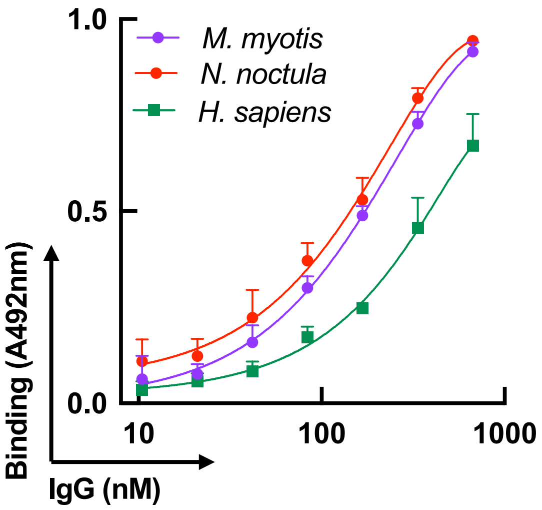


**Suppl. Figure 13.** Interaction of human and bat pooled IgG with hydrophobic hapten. Binding of increasing concentrations of IgG from *M. myotis* (purple line and symbols), *N. noctule* (red line and symbols) and human (green line and symbol) to immobilized protoporphyrin IX. Each data point on the graphs represents average optical density (n=4) ±SD from two independent experiments.
